# Axial Stress Provides a Lower Bound on Shear Wave Velocity in Active and Passive Muscle

**DOI:** 10.1101/2021.12.04.471223

**Authors:** Michel Bernabei, Sabrina S. M. Lee, Eric J. Perreault, Thomas G. Sandercock

## Abstract

Ultrasound shear wave elastography can be used to characterize mechanical properties of unstressed tissue by measuring shear wave velocity (SWV), which increases with increasing tissue stiffness. Measurements of SWV have often been assumed to be directly related to the stiffness of muscle. Some have also used measures of SWV to estimate stress, since muscle stiffness and stress covary during active contractions. However, few have considered the direct influence of muscle stress on SWV, independent of the stress-dependent changes in muscle stiffness, even though it is well known that stress alters shear wave propagation. The objective of this study was to determine how well the theoretical dependency of SWV on stress can account for measured changes of SWV in passive and active muscle. Data were collected from six isoflurane-anesthetized cats; three soleus muscles and three medial gastrocnemius muscles. Muscle stress and stiffness were measured directly along with SWV. Measurements were made across a range of passively and actively generated stresses, obtained by varying muscle length and activation, which was controlled by stimulating the sciatic nerve. Our results show that SWV depends primarily on the stress in a passively stretched muscle. In contrast, the SWV in active muscle is higher than would be predicted by considering only stress, presumably due to activation-dependent changes in muscle stiffness. Our results demonstrate that while SWV is sensitive to changes in muscle stress and activation, there is not a unique relationship between SWV and either of these quantities when considered in isolation.

## INTRODUCTION

Ultrasound shear wave elastography can be used to measure the mechanical properties of biological tissues non-invasively (1-4). In simple, homogeneous materials, shear wave velocity (SWV) provides an accurate estimate of the shear modulus and, through a direct relationship, the tensile or Young’s modulus, which is more relevant to how muscles act on the skeleton. The relatively low cost and ease of use of ultrasound systems for elastography has inspired many to use SWV to measure skeletal muscle properties (5, 6). However, skeletal muscle is a complex tissue and the relationship between its material properties and SWV is incompletely understood (7-9). This has led to SWV being used to estimate muscle force (10-12) and Young’s modulus (13-15), though it is unclear under which conditions either of these properties can be uniquely determined.

Muscle is unique in its ability to generate tension when activated – a process that leads to simultaneous changes in Young’s modulus that are thought to arise from increased cross-bridge attachment (16-19). The tensile load on a muscle changes with passive stretching, and these loads can also alter the Young’s modulus of muscle through the nonlinear stress-strain properties of its passive structures (20, 21). The change in muscle tension with activation and stretch complicates the relationship between SWV and material properties since it is well known that the speed with which a shear wave propagates changes with tension (22, 23), or more precisely stress. Traditional beam analysis shows that shear wave propagation depends on stress and material properties, specifically shear modulus (22, 24). Recently, Martin et al. (25) demonstrated that the analysis of shear wave propagation in beams provides a reasonable approximation for how they also propagate in tendons. In addition, they found that the speed with which a shear wave propagates along a tendon is primarily determined by stress. Muscle is obviously different from tendon. We previously demonstrated that SWV is sensitive to activation-dependent changes in muscle stiffness even when stress is held constant (26). We also found that stiffness alone could not account for the measured values of SWV but that stress must also be considered. Those results suggested that both stress and material properties contribute to the SWVs measured in muscle but precisely how they contribute and how their relative contributions change with passive stretching of muscle or activation remains unknown, preventing the use of SWV for uniquely determining either muscle stress or stiffness.

The objective of this study was to quantify how the SWV in skeletal muscle changes with passive lengthening and activation. Data were collected in the cat soleus and medial gastrocnemius muscles, allowing us to test the generalizability of our results across two muscles with very different fibre lengths and pennation angles. Our experimental preparation provided not only measures of SWV, but also simultaneous, direct measures of muscle stress and Young’s modulus. Experimental measures of SWV were compared to the stress-based model of SWV previously used to describe how SWV depends on the stress within tendons (25). This approach allowed us to test the hypothesis that a model of the relationship between stress and SWV can be used to characterize how shear waves propagate in muscle. This hypothesis was tested separately for passively stretched muscle and for muscle activated by electrical stimulation. Our results are the first to evaluate how a physics-based model of longitudinal stress influences SWV in living skeletal muscle. They demonstrate the utility of this simple characterization for passively stretched muscle and the magnitude of its limitations when muscle is active. Together, these results clarify the information that can be attained from shear wave elastography during the passive and active conditions altering force and stiffness in living muscles.

## METHODS

### Animal preparation

The purpose of this study was to determine if the stress alone can be used to estimate SWV in muscles with different architectures during active and passive conditions. We chose the cat soleus and cat medial gastrocnemius (MG) as our test muscles. Soleus is a slow fatigue resistant muscle with low pennation angle (about 7°) at muscle lengths producing peak tetanic tension (*L*_*o*_) and relatively long fascicles (42 mm at *L*_*o*_). In contrast MG is fast fatigable muscle, quite pennate (21° at *L*_*o*_) and with short muscle fibres (21 mm at *L*_*o*_). We measured SWV and stress simultaneously, testing the two muscles at different lengths and different levels of activation to obtain a range of passive and active stresses. Because we previously demonstrated that SWV varies with muscle stiffness (26) we also measured short range stiffness to determine if it can explain changes in SWV that could not be accounted for by the stress model alone.

We measured data from six female cats (Felis catus; 4.3 to 5.0 kg). Three for soleus and three for the MG. All animals were lawfully acquired from a designated breeding establishment for scientific research, and housed at Northwestern University’s Center for Comparative Medicine, an AAALAC accredited animal research program. The Institutional Animal Care and Use Committee at Northwestern University approved all procedures.

The muscle under study, either the soleus or MG, was isolated by partially removing the skin and the connective tissues of the crural fascia and then carefully dissecting the triceps surae muscle-tendon unit. We preserved the nerve and blood supply to the test muscle. Other muscles were denervated by cutting the peroneus nerve, the tibial nerve distal to the branch to soleus or MG, and the high muscular branch of the sciatic nerve. The distal tendon of soleus or MG was dissected free from the rest of the Achilles tendon. A bone chip was left attached to the tendon to provide a secure attachment, and secured to a servomechanism (Thrust tube linear motor; Dunkermotoren GmbH, Bonndorf, Germany) that allowed to generate fast, controlled changes in length. Force on the muscle-tendon unit force was measured with a load cell (Sensotec model 31; Honeywell, Golden Valley) mounted on the motor shaft. The knee and ankle joints were fixed to metal clamps. The whole left hindlimb was submerged in a temperature controlled saline bath (26°C) to permit ultrasound imaging without the transducer contacting the muscle. For a figure and more detailed information about the recording setup see (26).

### Nerve stimulation

A bipolar cuff electrode was folded around the sciatic nerve. The electrode was connected to a stimulus isolation unit (Grass stimulator S8800) that was under computer control. We generally used supramaximal currents in our stimulus pulses except for a few low force trials where we deliberately stimulated the muscle at sub-maximal levels. One twitch was evoked before each contraction to enable the muscle to adapt to its current length, minimizing history effects. Five-second trains were needed to allow muscle force to stabilize before measuring SWV. Such long trains created a problem with fatigue. In the soleus we found that 100 Hz for 5s produced substantial high frequency fatigue. This fatigue recovered quickly with rest but still created a 20% change in force within a trial (27). This was avoided in the soleus by using 40 Hz trains, which were still sufficient to maintain tetanic contractions. In the MG fatigue could not be avoided. In a single trial, force often fell by 30% from the start of the train until the end. A synchronization pulse from the ultrasound unit allowed the SWV measurements to be associated with the correct force. A 2-min recovery time was allowed between subsequent stimulations. However, even with this rest, force from the MG continued to fall throughout the experiment. This was not a major problem since our goal is to compare stress with SWV. We did not see any evidence that fatigue changed the relationship between force and SWV.

### Short Range Stiffness

A direct measure of muscle stiffness can be obtained by characterizing short-range stiffness (SRS). SRS describes the elastic properties of muscle in response to small, rapid changes in length. Measurements of SRS can be made in an in situ muscle preparation that allows for direct measures of muscle force and length. The SRS of individual muscles scales linearly with actively generated force and is thought to be directly related to the number of attached cross-bridges. We measured SRS with a fast length perturbation before the end of tetanic stimulation (26). The perturbation had a displacement of 2 mm and a speed of 2 m/s. For the cat soleus, this corresponded to a displacement of ∼2.7% of the soleus fascicle length at *L*_*o*_ (*x*_*fo*_) and a nominal speed of 54 *x*_*fo*_ /s. For cat MG the displacement was 5.4% of fascicle length with a speed of 108 *x*_*fo*_ /s. SRS is nearly invariant to the speed of stretch within this range (28).

### Shear wave ultrasound

An Aixplorer V9.1.1 ultrasound system (Supersonic Imagine, Aix-en-Provence, France) coupled with a linear transducer array (4–15 MHz, 256 elements, SuperLinear 15-4; Vermon, Tours, France) was used for all ultrasound elastography. The parameters of the system were set to 1) mode: MSK—foot and ankle, 2) opt: std, 3) persist: no, to maximize the sampling rate and avoid bias due to locked proprietary processing. The research grade software on our system provides shear wave motion images sampled at 8 kHz in sequences of 42 frames for each acoustic radiation pulse, recorded simultaneously with a single B-mode ultrasound image. The size of the region of interest was selected to include the belly of the muscle (up to 4×1 cm), and the speed of the compressional sound wave in the tissues was assumed to be 1,540 m/s. A custom-built frame held the transducer in a constant position relative to muscle, with the transducer oriented parallel to the fascicle plane.

### Data collection

A MATLAB XPC controller (MathWorks, Natick, MA) was used to control the muscle puller, sequence nerve stimulation, and record all data. Acquisition of ultrasound images and collection of the muscle data were synchronized with a trigger pulse from the ultrasound machine.

The experiment began by measuring the forces needed to accelerate the puller shaft, membrane, and saline in the bath, using the same step change in displacement used to measure SRS. This was later subtracted from measurements made on the muscle. Next, we measured the length-tension properties of the muscle. This set the operating range for the puller. We always kept the muscle at *L*_*o*_ minus 5mm during rest periods to keep passive force near zero. This minimized the stretch of the tendon-aponeurosis. We then began data collection. The muscle was moved to the test length prior to each 5 s trial. First a passive trial was measured, followed two seconds later an active trial; this time was sufficient for repositioning the probe as needed to optimize image acquisition during muscle stimulation. One B-mode image and one shear wave movie were captured simultaneously for each passive and active condition. The muscle was moved back to rest length for two minutes and the process repeated. We measured the force-SRS-SWV properties at three different lengths: i) *medium* (*L*_*o*_) to get the maximum active force, ii) *short*, a length chosen so that active force was about half of the peak force, iii) *long*, where active force was again half the peak value.

At the long length there was substantial passive tension. See Fig 1A for the length tension curve and measured force in the soleus. Most active measurements were made using supramaximal stimulation, but we also used sub-maximal stimulation in some measurements made at *L*_*o*_. This was to obtain multiple levels of active force at the same level of passive stretch. Finally, we measured passive muscle properties at long lengths. These were conducted at the end of each experiment as the imposed stretch exceeded the physiological length of muscle, eventually resulting in muscle damage. We found that the physical limits of the muscle were typically reached when passive force was equal to the peak active force.

**Fig 1).**
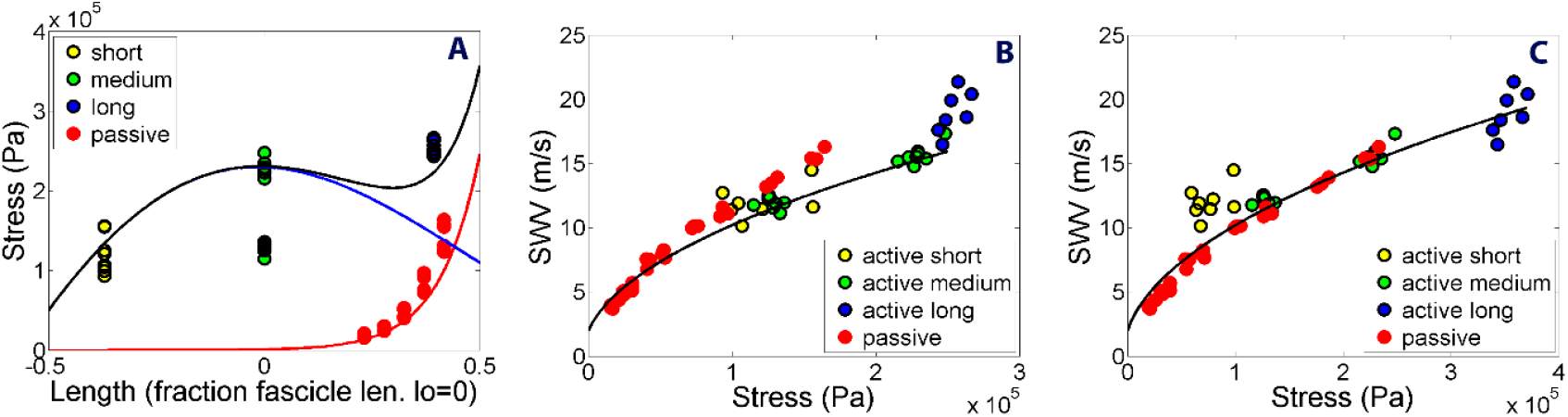
Typical data from cat soleus muscle. The same data points are displayed in each of the 3 plots. Red circles denote passive tension; Yellow, green and blue circles denote active muscle at respective muscle lengths of short, medium and long. A: shows the muscle lengths at which the data were measured. The solid red, blue, and black lines show passive, active, and total stress respectively. The green points, at *L*_*o*_ with half stress, were obtained using sub-maximal stimulation. B: SWV is plotted against stress. Here stress was calculated using the cross-sectional area at *L*_*o*_. The solid black line, here and also in plot C, denote the predicted results from the stress model (eq. 1). C: SWV is again plotted against force but here force is scaled by area varied as a function of muscle length. See text for details.

Positioning the ultrasound probe during active trials was difficult, particularly for the MG in which fascicles shorten by about 5mm during a tetanic contraction, increasing the angle of pennation (29). By trial-and-error we were able to position the probe to obtain clear images. However, this trial-and-error process led to increased fatigue and tendon stretch while the optimal position was determined.

### Data Processing

In this manuscript we present a simplified model of shear wave propagation, which we refer to as the stress model. We assume muscle is a homogenous, anisotropic, infinitely large material. The model considers a plane wave propagating along the muscle fibres with minimal contribution of the bending moment. If we also assume that the transverse shear modulus of the muscle did not change with stress, the constitutive equation for wave propagation derived from Timoshenko beam theory can be reduced to (25):

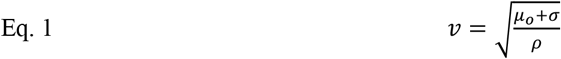

In this equation, *v* is the SWV along the muscle fibres in m/s, *µ*_*o*_ is the shear modulus of unstressed passive muscle in Pa, *σ* is stress (tension per unit area) in Pa, and *ρ* is tissue density in kg/m^3^.

SWV was estimated offline using displacement movies created by the Aixplorer ultrasound machine, as we detailed previously (26), and summarized here. Displacement of the wave was displayed as a function of time, depth, and distance along the fascicle. There were 5 separate pushes covering the selected area of the movie. The B mode image was used to further select part of the displacement movie that covered only the muscle of interest without any aponeuroses. The displacement movie was then averaged over this depth to create a 2D image where displacement is a function of time and distance along the fascicle. For each push the 2D displacement image was fit with a regression line the slope of which was an estimate of the SWV.

We used muscle cross sectional area to compute stress from the measured experimental forces. Cross sectional area at *L*_*o*_ was estimated using the maximum tetanic tension at *L*_*o*_, assuming the specific tension for muscle is 230 kPa (30, 31).

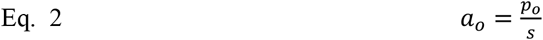

where *a*_*o*_ is the cross-sectional area at *L*_*o*_, *p*_*o*_ is tetanic tension at *L*_*o*_, and *s* is the specific tension.

Muscle cross sectional area is not constant but rather changes with passive stretching and activation. The influence of both can be approximated by considering the change in muscle fiber length relative to *L*_*o*_ such that the area at any length is given by:

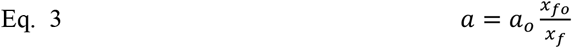

Mean fascicle length was estimated using muscle weight and density (1060 kg/m^3^) to estimate volume, and dividing by area to determine mean fascicle length at *L*_*o*_, *x*_*fo*_:

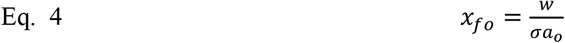

where *w* is the weight of the muscle, and σ is the density.

When a muscle was passively stretched to long lengths, it produced substantial tension. For example, the MG, stretched to a force equal to *p*_*o*_, causes a tendon-aponeurosis stretch of about 5 mm (29, 32). This tendon stretch was estimated and used to correct estimates of fascicle length. The assumed stretch was:

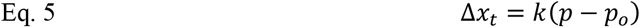

where *Δx*_*t*_ is change in tendon length due to changing force, *k* is the stiffness of the tendon-aponeurosis, *p* is the current force and *p*_*o*_ is the tetanic force at *L*_*o*_. The stiffness of the tendon-aponeurosis, *k*, was estimated using the model of Cui et al. (30).

This approach allowed us to compute the fascicle length, *x*_*f*_, as:

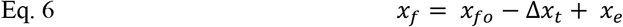

where *x*_*e*_ is the experimentally measured change in muscle-tendon length from *L*_*o*_.

The distributions of errors resulting from the stress model (Eq. 1) were analysed using a signed-rank test to determine if the mean for active and passive conditions differed from zero. Linear mixed effects models were used to describe these errors as functions of either short range stiffness or muscle stress. In both models, cat was considered as a random factor. Analysis was conducted in MATLAB 2019a. All results are presented as mean ± standard error, unless noted.

## RESULTS

The anatomical parameters from all muscles used in the study are shown in table 1.

**Table 1).** Anatomical parameters

| Muscle | $p_o$ (N) | Weight (g) | Area (cm<sup>2</sup>) | fascicle length (mm) | k tendon-aponeurosis (N/mm) |
| --- | --- | --- | --- | --- | --- |
| soleus 14 | 22 | 3.9 | 0.96 | 43.2 | 13.4 |
| soleus 15 | 27 | 4.3 | 1.17 | 38.8 | 18.9 |
| soleus 16 | 27 | 4.8 | 1.17 | 43.3 | 14.5 |
| MG 18 | 143 | 14.9 | 6.22 | 25.4 | 32 |
| MG 19 | 140 | 16.4 | 6.09 | 28.6 | 37 |
| MG 20 | 146 | 16.6 | 6.35 | 27.6 | 31 |

A typical example of SWV versus stress in the cat soleus is shown in Fig. 1. Fig. 1A shows the relation between the conditions imposed in the experiment and the muscle length-tension curve. The largest active stresses (about 230 kPa) were obtained at *L*_*o*_ (0 mm). Using sub-maximal stimulus currents, lower stresses (about 120 kPa) were measured at *L*_*o*_ (green points at lower stress). Note that at this length passive stress is near zero. By shortening the muscle, we could stimulate maximally and obtain less stress (yellow points). Stretching the muscle to long lengths and fully activating it also produced less active stress but significant passive stress (blue points). The red points show data when the muscle was passively stretched. Note that this involved some unusually long lengths beyond physiological limits. Fig. 1B shows the same data set with SWV plotted as a function of stress. Force was normalized by the fixed area *a*_*o*_ (Eq. 2). The solid black line represents the stress model. Fig. 1C shows the same data but now normalized by length-dependent calculations of area, estimated using fascicle length (*a* in Eq. 3). Normalization has the greatest influence on points measured at long lengths. In this animal, active and passive points measured at long lengths were similar to the stress model. Active short length data points lie above the stress model.

Passive data from all cats are shown in Fig 2A. The SWV for passively stretched muscle is described well by the stress model when the estimate of cross-sectional area is corrected to account for changes at different muscle lengths. SWV measurements from both the MG and soleus are close to what is predicted by the stress model.

**Fig 2).**
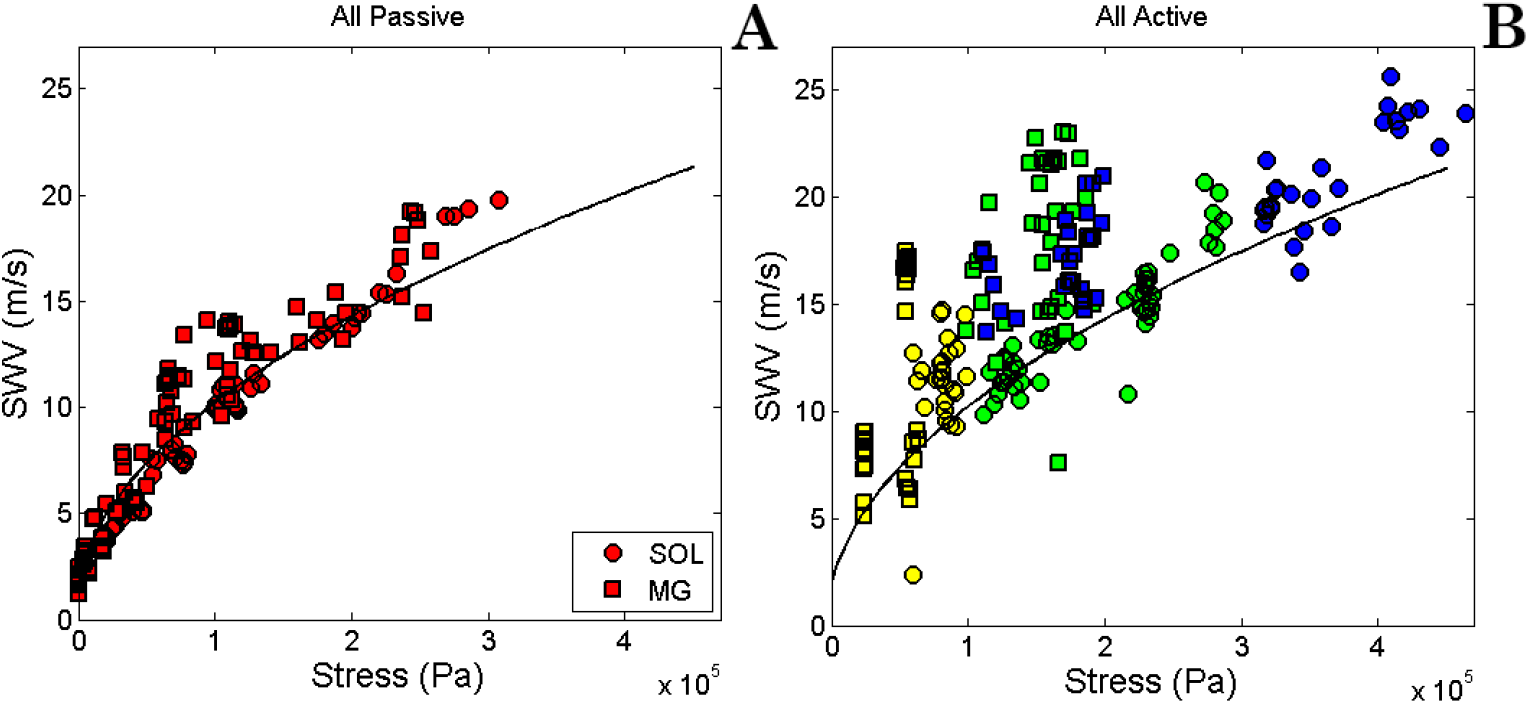
Data from all cats. Circles denote data from the soleus, squares from the MG muscles. Black lines denote the predicted results from the stress model (eq. 1) Panel 2A shows passive results. Panel 2B shows the active data. Yellow, green and blue show total force at short, medium, and long lengths.

The active trials from all cats are shown in Fig 2B. Note that stresses for some measurements are very high (∼2 times *p*_*o*_), probably higher than the physiological range of the muscle. This is due to the measurements being obtained at long lengths. Fig. 3 shows the same data, but for each cat shown separately. All 6 muscles show some active points that are substantially above the black line from the stress model. Note that the active trials in MG show large variability, likely due to the large change in pennation angle that occurs with MG activation and the challenges that poses for ultrasound measurements.

**Fig 3).**
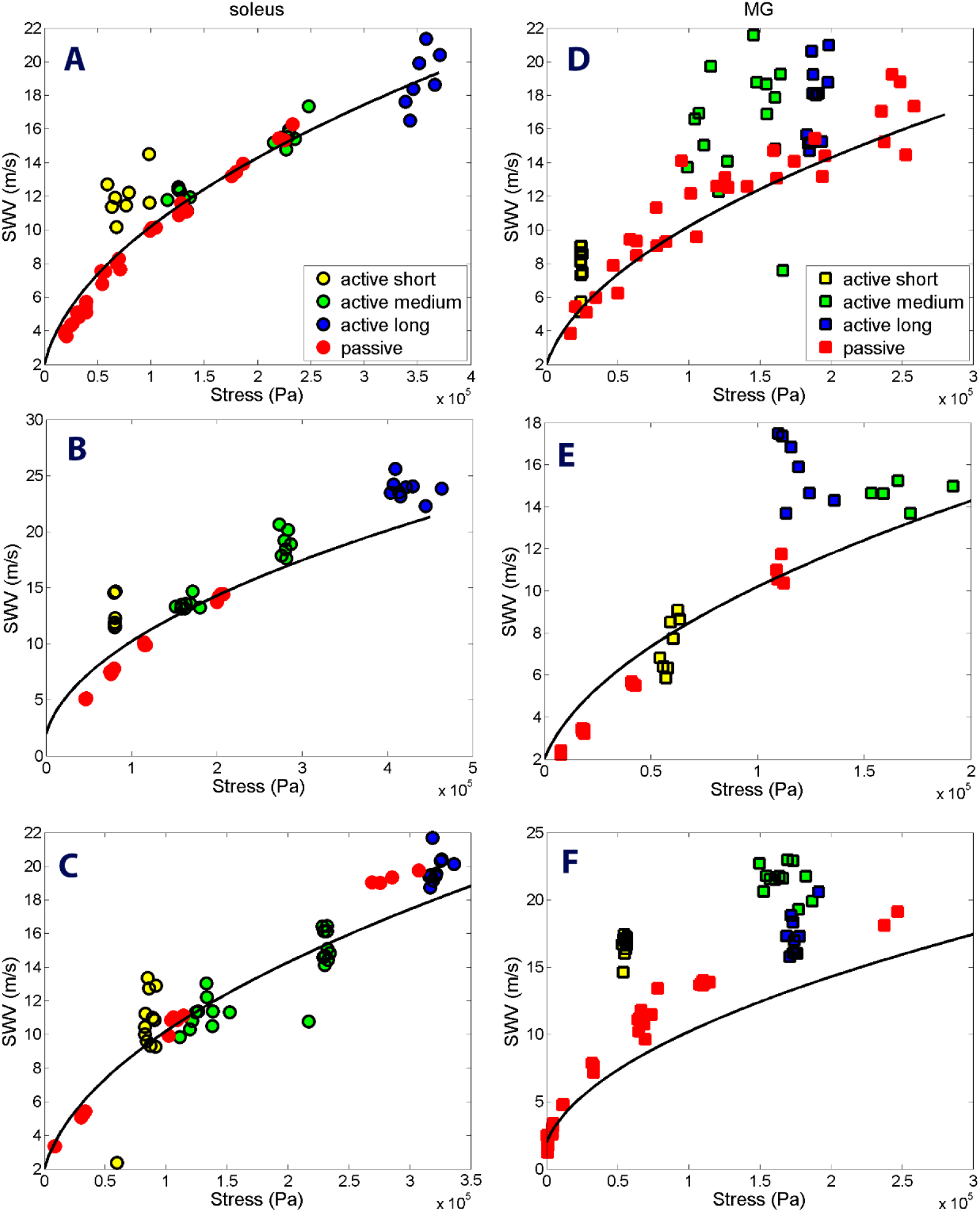
Similar to figure 2 except results from each cat are plotted independently. The left column is from soleus. The right column from MG.

The error in SWV, that is the experimental value minus the prediction from the stress model, is shown in Fig 4. Fig. 4A shows the distribution or errors for passive (mean=0.20 kPa, sdev=1.42) and active conditions (mean=2.72 kPa, sdev=3.12). A Kolmogorov-Smirnov test (kstest, MATLAB) showed that the distribution of active errors was non-Gaussian (p<.00001). We therefore used a Wilcoxon signed-rank test (signrank, MATLAB) to compare the mean of each distribution to zero. We found that the mean of the passive distribution was not significantly different from zero (p=.99), but that the mean of the active distribution was different (p<.000001). Because the errors were significantly different from zero for active muscles, we attempted to model these residual errors as a function of both muscle stress and the independently measured short-range stiffness (SRS). Fig. 4B shows the error plotted as a function of stress for both active and passive conditions. Fig. 4C shows the same data plotted against SRS. In both cases, data from all animals are plotted.

**Fig 4).**
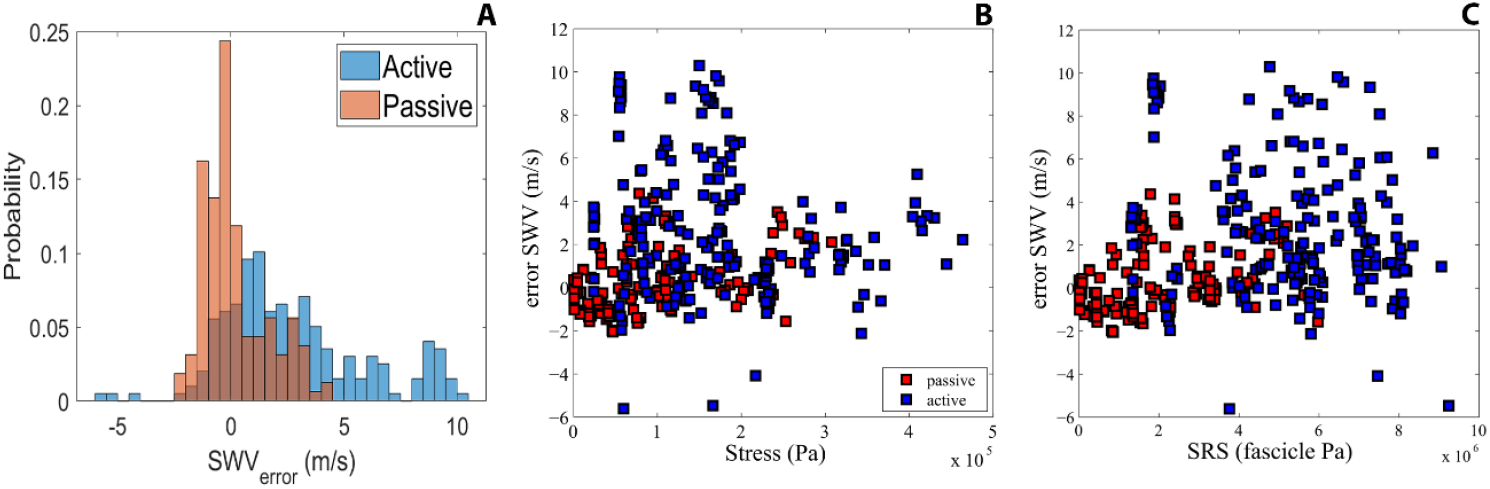
SWV errors. All SWV measurements have had the stress model subtracted. Red shows passive, blue total tension. A) Distribution of errors. Passive error is close to a gaussian distribution. Active error has a strong bias toward high SWV. B) error plotted against tension. B) error plotted against SRS of fascicle. See text for statistics.

The error in SWV was correlated with our measures of SRS in both active and passive muscle. This was assessed using a linear mixed effects model with SRS and activation state of the muscle (passive or active) as the independent variables and the error difference between the squared value of the measured SWV (SWV^2^) and that predicted by the stress model as the dependent variable (Fig. 5). SWV^2^ was used to yield a linear relationship between our independent and dependent variables (Eq. 1). Cat was considered as a random factor having an independent effect on the average slope and intercept across activation states. The error in the SWV^2^ increased significantly with SRS for both passive (p=0.0007) and active (p=0.0011) conditions, though there was no significant difference in these slopes (p=0.47). The errors in the active conditions were offset from those in the passive conditions (Δ=48±21 m^2^/s^2^; p=0.02). The R^2^ for the overall model was 0.64 indicating that there remains a significant amount of measurement variance not accounted for by SRS.

**Fig. 5).**
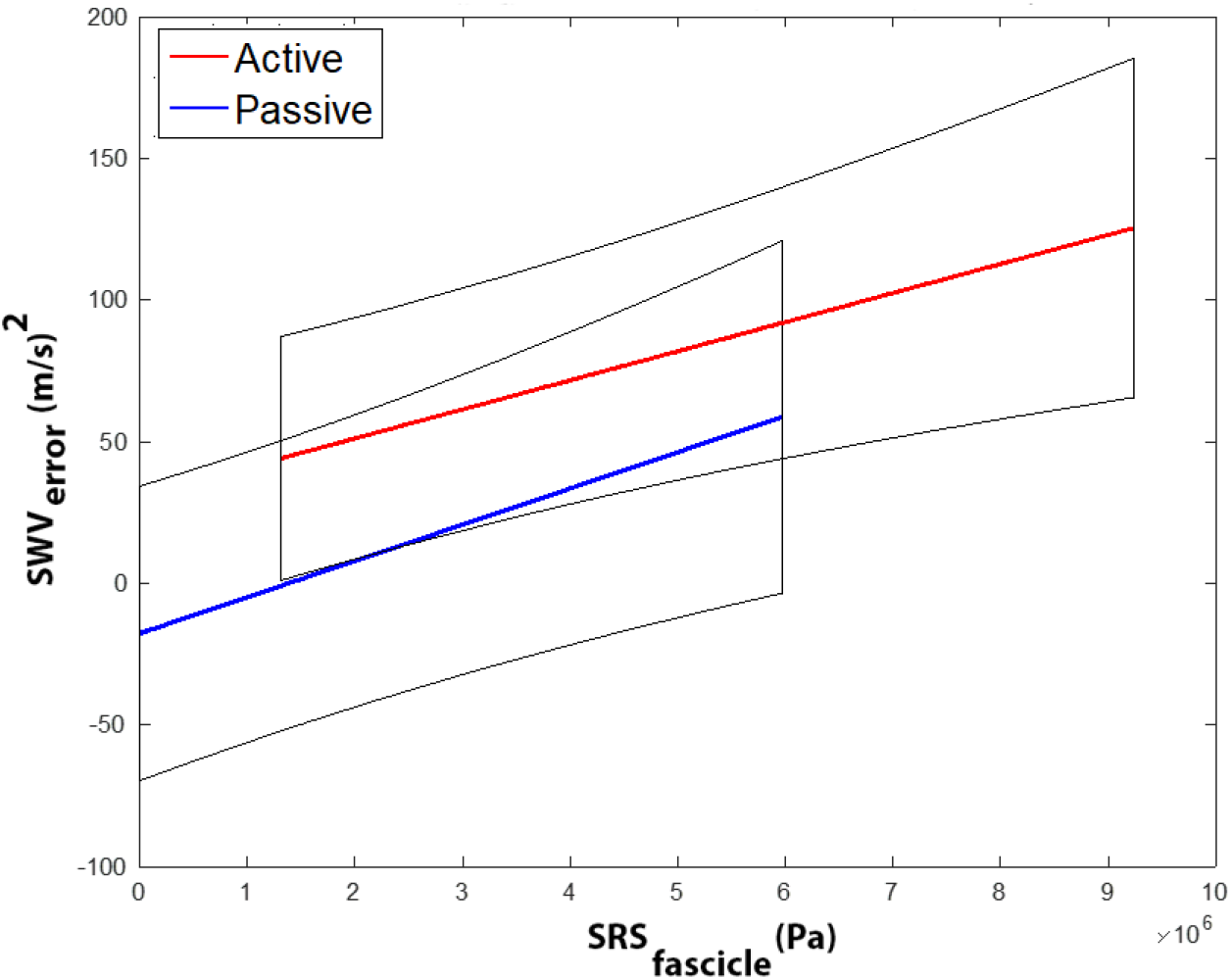
Dependence of stress model error on muscle short range stiffness (SRS). This relationship was assessed using a linear mixed effects model with SRS and activation state of the muscle (passive or active) as the independent variables and the error difference between the measured SWV2 that predicted by the stress model as the dependent variable. Cat was considered as a random factor having an independent effect on the average slope and intercept across activation states. A linear mixed effects model was used to determine if SRS accounted for some of the error. SWV error squared was treated as the dependent variable, SRS as continuous independent factor, and cat as a random factor.

SRS was strongly correlated with muscle stress in each animal, having an average R^2^ of 0.97±0.01 for the passive conditions and 0.70±0.05 for the active conditions. For these reasons modeling the SWV^2^ as a function of stress (R^2^=0.63) worked almost as well as when it was modeled as a function of stiffness. However, when both stress and SRS were considered as independent variables, only SRS had a significant effect on the error in SWV^2^ (SRS: p<0.0001; stress: p=0.3).

## DISCUSSION

The aim of this study was to examine the relationship between muscle stress and SWV in passive and active muscle tissues. We applied controlled changes in force by lengthening and activating the soleus and MG muscles of the cat. Our primary hypothesis was that changes in SWV can be predicted by a simple model accounting for muscle stress and knowledge of the SWV for unstressed muscle. Our results show that this stress model fits passive data well but only sets the lower bound for SWV measured in active muscle. Active muscle showed more variability and often had SWV values well above what was predicted by the stress model. These experiments were not designed to determine if the increased SWV was due to increased muscle stiffness, but activation dependent changes in stiffness appears to account for some of the error.

We are the first to show that SWV in passive muscle can be predicted by stress. This is important because stress and stiffness covary in muscles. It has proven difficult to decouple the effect of one from the other on human muscle tissue because only indirect measures of force or stiffness are possible (7, 33). Our results are in line with a recent study on tendon tissues where Martin et al. (25) demonstrated that tensional pre-load is the main determinant of shear wave propagation in tendon, with the shear modulus playing a role only at the lowest tensions. Other studies measured SWV in animal muscle ex-vivo (8, 34), showing SWV increased with increasing tension. Eby et al. also measured Young’s modulus, showing it increased with stress. They suggested Young’s modulus accounted for the change in SWV. We believe their observations can, in part, be attributed directly to tension rather than Young’s modulus. Our analysis of Eby et al.’s data show the stress model accounts for about half of their measured SWV. The difference between our results and theirs may be due to our use of living muscle tissue, whereas Eby et al’s studied excised, dead tissue, which might have a higher stiffness that contributed to their higher SWV for a given stress. Still, the Eby results are consistent with our main conclusion, which is that the stress model sets a lower bound on the SWV within muscle tissue.

Our data show the stress model is incomplete for active muscle. It still appears to set a lower bound but underestimates measures SWV versus stress. Others have shown SWV increases in active muscle, but did not have a direct measure of muscle stress (33, 35, 36). So, we are unaware of direct comparisons that can be made to our experimental results. Understanding shear wave propagation in contracting muscle presents a challenge because active muscle tension and SRS both depend on the number of attached cross-bridges. In a previous attempt to clarify the active stress-stiffness-SWV relationship, we demonstrated that SWV varies with muscle stiffness when tension is held constant (26). We manipulated muscle temperature to elicit changes in muscle stiffness (SRS) without changes in active force. Here, we show that SRS explains part of the error in the stress model, emphasizing the importance of understanding how both muscle stress and stiffness contributed to SWV in active muscle.

There are some limitations to our study. We had a surprising amount of variability in our data, particularly for active measurements in the MG. We have not identified the source, but can formulate several hypotheses. First, we expect greater variance in SWV at high velocities because of limited sampling time, but there is no reason to believe the error should be greater than data obtained from passive muscle at the same tension. Second, positioning the ultrasound probe was more difficult in active muscle because of movement of the muscle relative to the probe. This was particularly true for the MG, which had substantial changes in pennation angle during active contractions potentially explaining the increased variability in this muscle. However, it should not lead to an overall increase in SWV compared to passive muscle or the stress model. A third problem seems to be the quality of the displacement movie used to calculate SWV. Our subjective impression was that the clarity of the B-mode images, and displacement movie, were not as good in active muscle, particularly at short muscle lengths. The fascicles did not always stand out as clearly. Combined with the heterogeneity of active stress within the muscle at short lengths or sub-maximal activation, this could be another reason we see so much variability.

In summary, we found that a theoretical model of how stress influences SWV predicted our experimental results well for passively stretched muscles. In contrast, this same model significantly underpredicted the SWV in active muscles. These findings were consistent in the soleus and medial gastrocnemius muscles, suggesting our results generalize across muscles with different architectures and fiber types. These findings demonstrate that SWV is sensitive to changes in muscle stress and that a model of how stress influences SWV can be used to predict the SWV in passively stretched living muscles. This same model provides only a lower bound on the SWV in active muscle, presumably due to activation-dependent changes in muscle stiffness. Together, our results provide further clarity on the factors influencing shear wave propagation in muscle.

## Acknowledgments

Supported by National Institute of Health Grant 5R01-AR-071162-02 to E. J. Perreault.

## Author contributions

M.B., S.S.M.L., E.J.P., and T.G.S. conceived and designed research;

M.B. and T.G.S performed experiments; M.B., T.G.S., and E.J.P. analyzed data;

M.B., S.S.M.L., E.J.P., and T.G.S. interpreted results of experiments;

T.G.S prepared figures;

M.B. and T.G.S drafted manuscript;

M.B., S.S.M.L., E.J.P., and T.G.S. 378 edited and revised manuscript;

M.B., S.S.M.L., E.J.P., and T.G.S. approved final version of manuscript

## REFERENCES

1. Bercoff J, Tanter M, and Fink M. Supersonic shear imaging: a new technique for soft tissue elasticity mapping. IEEE Trans Ultrason Ferroelectr Freq Control 51: 396–409, 2004.

2. Cosgrove DO, Berg WA, Doré CJ, Skyba DM, Henry JP, Gay J, and Cohen-Bacrie C. Shear wave elastography for breast masses is highly reproducible. European radiology 22: 1023–1032, 2012.

3. Frulio N, and Trillaud H. Ultrasound elastography in liver. Diagnostic and interventional imaging 94: 515–534, 2013.

4. Gennisson JL, Deffieux T, Macé E, Montaldo G, Fink M, and Tanter M. Viscoelastic and anisotropic mechanical properties of in vivo muscle tissue assessed by supersonic shear imaging. Ultrasound in medicine & biology 36: 789–801, 2010.

5. Gennisson JL, Deffieux T, Fink M, and Tanter M. Ultrasound elastography: principles and techniques. Diagnostic and interventional imaging 94: 487–495, 2013.

6. Murayama M, Nosaka K, Yoneda T, and Minamitani K. Changes in hardness of the human elbow flexor muscles after eccentric exercise. European journal of applied physiology 82: 361–367, 2000.

7. Chernak LA, DeWall RJ, Lee KS, and Thelen DG. Length and activation dependent variations in muscle shear wave speed. Physiological measurement 34: 713–721, 2013.

8. Eby SF, Song P, Chen S, Chen Q, Greenleaf JF, and An KN. Validation of shear wave elastography in skeletal muscle. Journal of biomechanics 46: 2381–2387, 2013.

9. Royer D, Gennisson JL, Deffieux T, and Tanter M. On the elasticity of transverse isotropic soft tissues (L). The Journal of the Acoustical Society of America 129: 2757–2760, 2011.

10. Killian B, Nordez A, and Hug F. Estimation of Individual Muscle Force Using Elastography. PLOS ONE 6: e29261, 2011.

11. Koo TK, Guo JY, Cohen JH, and Parker KJ. Quantifying the passive stretching response of human tibialis anterior muscle using shear wave elastography. Clin Biomech (Bristol, Avon) 29: 33–39, 2014.

12. Bouillard K, Hug F, Guével A, and Nordez A. Shear elastic modulus can be used to estimate an index of individual muscle force during a submaximal isometric fatiguing contraction. Journal of applied physiology (Bethesda, Md : 1985) 113: 1353–1361, 2012.

13. Bensamoun SF, Ringleb SI, Littrell L, Chen Q, Brennan M, Ehman RL, and An KN. Determination of thigh muscle stiffness using magnetic resonance elastography. J Magn Reson Imaging 23: 242–247, 2006.

14. Deffieux T, Montaldo G, Tanter M, and Fink M. Shear wave spectroscopy for in vivo quantification of human soft tissues visco-elasticity. IEEE Trans Med Imaging 28: 313–322, 2009.

15. Maïsetti O, Hug F, Bouillard K, and Nordez A. Characterization of passive elastic properties of the human medial gastrocnemius muscle belly using supersonic shear imaging. Journal of biomechanics 45: 978–984, 2012.

16. Proske U, and Morgan DL. Stiffness of cat soleus muscle and tendon during activation of part of muscle. Journal of neurophysiology 52: 459–468, 1984.

17. Rack PM, and Westbury DR. The short range stiffness of active mammalian muscle and its effect on mechanical properties. J Physiol 240: 331–350, 1974.

18. Ettema GJ, and Huijing PA. Skeletal muscle stiffness in static and dynamic contractions. Journal of biomechanics 27: 1361–1368, 1994.

19. Morgan DL. Separation of active and passive components of short-range stiffness of muscle. 951201 232: C45–49, 1977.

20. Roberts TJ. Contribution of elastic tissues to the mechanics and energetics of muscle function during movement. The Journal of experimental biology 219: 266–275, 2016.

21. Lieber RL, and Binder-Markey BI. Biochemical and structural basis of the passive mechanical properties of whole skeletal muscle. J Physiol 599: 3809–3823, 2021.

22. Graff KF. Wave Motion in Elastic Solids. Dover Books, 1991.

23. Resnick R, Halliday D, and Krane KS. Physics, Volume 1 5th Edition. John Wiley and Sons, 2002.

24. Remeniéras JP, Bulot M, Gennisson JL, Patat F, Destrade M, and Bacle G. Acousto-elasticity of transversely isotropic incompressible soft tissues: characterization of skeletal striated muscle. Physics in medicine and biology 66: 2021.

25. Martin JA, Brandon SCE, Keuler EM, Hermus JR, Ehlers AC, Segalman DJ, Allen MS, and Thelen DG. Gauging force by tapping tendons. Nature Communications 9: 1592, 2018.

26. Bernabei M, Lee SSM, Perreault EJ, and Sandercock TG. Shear wave velocity is sensitive to changes in muscle stiffness that occur independently from changes in force. Journal of applied physiology (Bethesda, Md : 1985) 128: 8–16, 2020.

27. Sandercock TG, Faulkner JA, Albers JW, and Abbrecht PH. Single motor unit and fiber action potentials during fatigue. Journal of applied physiology (Bethesda, Md : 1985) 58: 1073–1079, 1985.

28. Cui L, Perreault EJ, and Sandercock TG. Motor unit composition has little effect on the short-range stiffness of feline medial gastrocnemius muscle. Journal of applied physiology (Bethesda, Md : 1985) 103: 796–802, 2007.

29. Griffiths RI. Shortening of muscle fibres during stretch of the active cat medial gastrocnemius muscle: the role of tendon compliance. 951201 436: 219–236, 1991.

30. Cui L, Perreault EJ, Maas H, and Sandercock TG. Modeling short-range stiffness of feline lower hindlimb muscles. Journal of biomechanics 41: 1945–1952, 2008.

31. Sacks RD, and Roy RR. Architecture of the hind limb muscles of cats: functional significance. Journal of Morphology 173: 185–195, 1982.

32. Griffiths RI. The mechanics of medial gastrocnemius muscle in the freely hopping wallaby (Thylogale billardierii). 951201 147: 439–456, 1989.

33. Bouillard K, Nordez A, and Hug F. Estimation of individual muscle force using elastography. PLoS One 6: e29261, 2011.

34. Koo TK, Guo JY, Cohen JH, and Parker KJ. Relationship between shear elastic modulus and passive muscle force: an ex-vivo study. Journal of biomechanics 46: 2053–2059, 2013.

35. Hug F, Tucker K, Gennisson JL, Tanter M, and Nordez A. Elastography for Muscle Biomechanics: Toward the Estimation of Individual Muscle Force. Exercise and sport sciences reviews 43: 125–133, 2015.

36. Nordez A, and Hug F. Muscle shear elastic modulus measured using supersonic shear imaging is highly related to muscle activity level. Journal of applied physiology (Bethesda, Md : 1985) 108: 1389–1394, 2010.

